# Chemogenetically decreasing activity of the dorsal cochlear nucleus can ameliorate noise-induced tinnitus in mice

**DOI:** 10.1101/2021.04.06.438667

**Authors:** Thawann Malfatti, Barbara Ciralli, Markus M. Hilscher, Richardson N. Leao, Katarina E. Leao

## Abstract

The dorsal cochlear nucleus (DCN) is a region known to integrate somatosensory and auditory inputs and is identified as a potential key structure in the generation of phantom sound perception, especially noise-induced tinnitus. Yet, how altered homeostatic plasticity of the DCN induces and maintains the sensation of tinnitus is not clear. Here, we chemogenetically decrease activity of a subgroup of DCN neurons, Ca^2+^/Calmodulin kinase 2*α* (CaMKII*α*) positive DCN neurons, using Gi-coupled human M4 Designer Receptors Exclusively Activated by Designer Drugs (hM4Di DREADDs), to investigate their role in noise-induced tinnitus. Mice were exposed to loud noise (9-11kHz, 90dBSPL, 1h, followed by 2h of silence) and auditory brainstem responses (ABRs) and gap prepulse inhibition of acoustic startle (GPIAS) were recorded two days before and two weeks after noise exposure to identify animals with a significantly decreased inhibition of startle, indicating tinnitus but without permanent hearing loss. Neuronal activity of CaMKII*α*+ neurons expressing hM4Di in the DCN was lowered by administration of clozapine-N-oxide (CNO). We found that acutely decreasing firing rate of CaMKII*α*+ DCN units decrease tinnitus-like responses (p = 0.038, n = 11 mice), compared to the control group that showed no improvement in GPIAS (control virus; CaMKII*α*-YFP + CNO, p = 0.696, n = 7 mice). Extracellular recordings confirmed CNO to decrease unit firing frequency of CaMKII*α*-hM4Di+ mice and alter best frequency and tuning width of response to sound. However, these effects were not seen if CNO had been previously administered during the noise exposure (n = 6 experimental and 6 control mice). Our results suggest that CaMKII*α*-hM4Di positive cells in the DCN are not crucial for tinnitus induction but play a significant role in maintaining tinnitus perception in mice.

## Introduction

Noise-induced tinnitus, commonly known as “ringing in the ears”, affects 10-15% of the world population (Heller, 2003; Gallus et al., 2015), where 1-2% seek medical assistance for severely decreased quality of life due to chronic tinnitus-related irritability, stress, anxiety and/or depression (Møller, 2007; Langguth et al., 2011; Shore et al., 2016). The origin of tinnitus pathophysiology have been linked to the dorsal cochlear nucleus (DCN) of the auditory brainstem (Kaltenbach et al., 2005; Tzounopoulos, 2008; Baizer et al., 2012; Shore et al., 2016; Shore and Wu, 2019), however, tinnitus generation and perception mechanisms are not well separated and far from completely understood.

Noise overexposure is known to alter firing properties of DCN cells (Brozoski et al., 2002; Finlayson and Kaltenbach, 2009; Pilati et al., 2012; Li et al., 2013; Manzoor et al., 2013), even after brief sound exposure at loud intensities (Gao et al., 2016). Such alterations within the DCN circuits could relay abnormal signaling to higher auditory areas and confound spontaneous firing with sensory evoked input, generating tinnitus. It has been suggested that noise-induced tinnitus is partly due to an imbalance of excitation and inhibition within the DCN (Kaltenbach and Manz, 2012; Shore et al., 2016) due to decrease in GABAergic (Middleton et al., 2011) and glycinergic activity (Wang et al., 2009) for example. On the contrary, excitatory fusiform cells have been shown to increase burst activity (Pilati et al., 2012; Wu et al., 2016) following noise overexposure. Furthermore, a shift in bimodal excitatory drive of the DCN after noise overexposure have been shown due to down-regulation of vesicular glutamate transport 1 (VGlut1; auditory-related) and up-regulation of VGlut2 (somatosensory related) proteins in the cochlear nucleus (Heeringa et al., 2018; Han et al., 2019). We have recently shown that directly manipulating activity of Ca^2+^/Calmodulin kinase 2*α* (CaMKII*α*) positive DCN neurons in vivo using optogenetics can have distinct effects on unit activity of the DCN, also in neurons not responding directly to neither sound or optogenetic light stimuli (Malfatti et al., 2021), highlighting how heavily interconnected the DCN circuit is (Oertel and Young, 2004). DCN circuit disruption such as bilateral electrolytic DCN lesioning in rats has shown to prevent tinnitus generation (Brozoski et al., 2011). Also, electrical stimulation of the DCN of rats can suppress tinnitus (Luo et al., 2012), and electrical high-frequency stimulation of the DCN with noise-induced tinnitus has shown to decrease tinnitus-perception during tests (van Zwieten et al., 2019). This indicates that unspecific alterations of DCN activity can decrease tinnitus induction and perception, but if the same DCN populations are involved in the two mechanisms remains to be investigated.

Here we behaviorally examine if tinnitus perception can be reduced by lowering the activity of CaMKII*α* positive DCN neurons using chemogenetics. We have recently shown this promoter to be expressed by both excitatory and inhibitory DCN neurons, but with a preference for slow-firing units (Malfatti et al., 2021), presumable excitatory fusiform cells (Ochiishi et al., 1998; Oh et al., 2014). We specifically investigated if noise-induced tinnitus, without hearing loss, can be ameliorated by lowering DCN neuronal activity. Next we decrease CaMKII*α*+ DCN neurons activity already during noise overexposure, to investigate if the same population is important for induction of tinnitus, and found that CaMKII*α*+ DCN neurons play different roles in induction and maintenance of noise-induced tinnitus.

## Methods

### Animals

Male C57Bl/6J mice (n=30) were used at the age of 21 days at first and 2 months at the last experiment, and were used for each step of the experimental timeline (see complete timeline in Figure 7A). All animal procedures were approved and followed the guidelines of the Ethical Committee of Animal Use (CEUA) from the Federal University of Rio Grande do Norte (CEUA protocol number 051/2015). Animals were housed on a 12h/12h day/night cycle and had free access to food and water.

### Gap prepulse inhibition of acoustic startle reflex

The gap prepulse inhibition of acoustic startle (GPIAS, Turner et al., 2006) test, based on the acoustic startle reflex in response to sudden loud sounds, was conducted in a sound-shielded room inside a sound-shielded chamber with LED lights. During recordings, the animal was placed inside a clear acrylic tube (Acrilart, Natal, Brasil), dimensions 6.1×5.9×5.1cm, with perforated plates closing the tube at both ends. The tube dimensions restricted mice from standing on the back paws. A speaker (Selenium Trio ST400, JBL by Harman, Brazil) was placed 4.5cm away from the restraining tube. In order to measure the animal’s startle reflex, a piezoelectric or a digital accelerometer was mounted to the base plate of the restraining tube. Sound stimulus consisted of blocks of narrow-band uniform white noise at background level, loud intensity (105dBSPL) or silence. Specifically, the stimulus was presented in the following sequence: a random integer value between 12 and 22 seconds of noise at background level (randomized background noise between trials); 40ms of noise at background level for Startle trials, or 40ms of silence for Gap-startle trials (Gap portion); 100ms of noise at background level (background noise before loud pulse); 50ms of noise at 105dBSPL (loud pulse); and 510ms of noise at background level (final background noise). Timestamp marks were used only for the loud pulse. The bands of frequencies tested were 8-10, 9-11, 10-12, 12-14, 14-16 and 8-18kHz. Background noise level was, for the initial GPIAS test, 60dBSPL. For GPIAS after noise exposure, background noise level was adjusted to 10dBSPL above the hearing threshold for the frequency tested.

Before each session the acrylic tube was cleaned with ethanol (70%) and next with water to remove residual smell of ethanol. Animals were habituated by handling for 10 minutes in the test room for two consecutive days followed by three days of acclimatization where animals were placed in the GPIAS tube and exposed to background noise, and next returned to their homecage. A successful acclimatization and habituation was considered when animals enter freely and do not urinate or defecate in the tube. After the habituation/acclimatization period, animals were screened for gap detection capability. The animals were placed in the restraining tube and left in the recording chamber for 5 minutes, allowing the animal to stay calm and stop exploring the chamber (Valsamis and Schmid, 2011). The test consisted of 18 trials per band of frequency tested, 9 with gap (Gap-startle trials) and 9 with noise filling the gap portion of the stimulus (Startle trials), presented pseudo-randomly. The GPIAS sessions were carried out at 3 time points for each animal. Initially, for screening animals before being included in experimental groups (see analysis for exclusion criteria), then in the end of the experiment timeline in the following NaCl injections, and the following day 30 min after CNO (0.5mg/kg, dissolved in dimethyl sulfoxide - DMSO at 3.3mg/ml, then diluted in NaCl to the final concentration of 50*μ*g/ml) administration. Each GPIAS session lasted between 23-41 min in total (depending on the randomization of inter pulse intervals). Upon the end of the session animals were returned to their home cage.

### Virus injection

Mice were anesthetized with an i.p. injection of ketaminexylazine combination at 90/6 mg/kg. When necessary, additional ketamine at 45 mg/kg was applied during surgery. The mouse was next mounted into a stereotaxic device resting on a heating block (37°C). The eyes were covered with dexpanthenol to prevent ocular dryness and povidone-iodine 10% was applied onto the skin of the animal’s head to avoid infections. The skin was anesthetized with lidocaine hydrochloride 3% before a straight incision was made, and hydrogen peroxide 3% was applied onto the exposed skull to remove connective tissue and visualize bone sutures. A small hole was carefully drilled at bilateral DCN coordinates (anteroposterior; AP=−6.24mm and mediolateral; ML=±2.3mm) using a dental microdrill. Next aliquoted virus (experimental: rAAV5/CaMKII*α*-HA-hM4D(Gi)-IRES-mCitrine, UNC Vector Core #AV4617C, viral concentration of 1.6×10^12^vm/ml; or control: rAAV5/CaMKII*α*-eYFP, UNC GTC Vector Core #AV4808D, 4.4×10^12^vm/ml) was rapidly thawed and withdrawn (1.5*µ*l) using a syringe pump (Chemyx NanoJet infusion pump). The needle (10*µ*l Nanofil syringe with a 34-gauge removable needle) was slowly inserted into the brain (dorsoventral; DV=4.3mm) and 0.75*µ*l of virus was infused (0.15*µ*l/min). At completed infusion, the needle was kept in the DV coordinate for five minutes to allow for the virus to diffuse, and then the needle tip was retracted to 3.8mm DV, where 0.75*µ*l of virus was again infused at the same rate. After the second infusion, the needle was kept in place for 10 minutes, to allow for a complete diffusion into the target area, before carefully removed. The same procedure was performed bilaterally. Following injections the skin was sutured, lidocaine hydrochloride 3% applied over the suture and 200*µ*l of NaCl subdermally injected for rehydration. Animals were monitored until fully recovered from anesthesia.

### Auditory brainstem responses

Similarly to the GPIAS setup, the speaker was connected to a sound amplifier connected to a sound card; and placed 4.5cm away from a stereotaxic frame. Field potentials (auditory brainstem responses - ABRs) were recorded using two chlorinated coiled Ag/AgCl electrodes as a recording and a reference electrode (1kΩ impedance). The electrodes were connected to the RHD2132 headstage through a DIP18-Omnetics connector, connected to Open-ephys board. Animals were anesthetized with an i.p. injection of ketamine-xylazine combination at 90/6 mg/kg and fitted to the stereotaxic frame, placed on an electric thermal pad and kept at 37°C. Dexpanthenol or NaCl was applied on the animal’s eyes to avoid drying of the ocular surface. Next the scalp was disinfected with polividone-iodine (10%) and two small incisions were made: one in the skin covering the lambda region and another in the skin over the bregma region. The electrodes were placed subdermally into the incisions and the ground was connected to the system ground. The electrode at bregma was used as reference, and the electrode over lambda was used for recording. Sound stimuli consisted of narrow-band uniform white noise pulses (3ms), presented at 10Hz for 529 repetitions for each frequency and intensity tested. The frequency bands tested were the same used for GPIAS: 8-10, 9-11, 10-12, 12-14 and 14-16kHz (with exception for the 8-18kHz frequency band); and sound pulses were presented at decreasing intensities from 80 to 35dBSPL, in 5dBSPL steps, with 10s of silence between different intensities. After the test, electrodes were removed, lidocaine hydrochloride 3% was applied on the incisions and 200*µ*l of NaCl was injected subdermally for rehydration. Animals were monitored after surgery until fully recovered from anesthesia and then returned to their home cage.

### Noise exposure

Anesthetized mice were placed inside a sound-shielded chamber, inside an acrylic tube, in an acoustically shielded room, with a speaker placed 4.5cm in front of the head of the mouse. Noise exposure consisted of narrow-band uniform white noise presented at 90dBSPL, 9-11kHz, for 1 hour. The animal was left in the acrylic tube, in the sound-shielded chamber for 2h following noise exposure, since external noise before and/or following noise exposure can interfere in tinnitus development (Norena and Eggermont, 2006; Sturm et al., 2017; Fan et al., 2020). During noise exposure and the silence period, the animal was monitored each 15 minutes and later returned to its homecage. Animals were given two days to recover before any further procedures. In some experiments CNO (0.5mg/kg) was given 30 min prior to noise exposure.

### *in vivo* unit recording

Animals were anesthetized with an i.p. injection of ketamine-xylazine combination at 90/6 mg/kg, and placed into the stereotaxic frame similar to for ABR recordings. A small craniotomy was drilled above the left DCN (AP=-6.24mm ML=−2.3mm) and a silicon depth probe (16 channels, 25 or 50*µ*m channel spacing, 177*µ*m recording site area, 5mm long shank; NeuroNexus A16) dipped in fluorescent dye (1,1’-dioctadecyl-3,3,3’,3’-tetramethylindocarbocyanine perchlorate; DiI, Invitrogen) for 10 minutes (for probe position) before lowered into the DCN (DV=4.3mm). A coiled Ag/AgCl wire soldered to a jumper wire was used as reference. The probe and reference wire were both connected to a headstage (RHD2132) through an adaptor (DIP18-Omnetics) connected to Openephys board, recording at a sampling rate of 30kHz. Sound stimulus consisted of narrow-band uniform white noise pulses (3ms) as described for ABRs, presented at 10Hz for 529 repetitions for each frequency and intensity tested. Spontaneous activity was recorded for 5 minutes, then the animal received an i.p. injection of NaCl, then sound stimulation started 30 minutes later. Subsequently, the same procedure was repeated for CNO (0.5mg/kg). At the end of the recording session the animals were either sacrificed by intracardial perfusion (20mL PBS and 20mL paraformaldehyde 4%) or by an overdose of ketamine followed by decapitation.

### Data analysis

All scripts used for controlling devices, stimulation control and data analysis are available online (LabScripts git repository, Malfatti, 2021a). The operating system of choice was Gentoo GNU/Linux, due to its flexible management of libraries (Ioanas, 2017). Recordings were done using Open-ephys GUI (Siegle et al., 2015). Microcontrollers and sound cards were controlled using SciScripts (Malfatti, 2021b), and the sounddevice python library (Geier, 2015) was used to read and write signals from/to the sound card. Calculations were done using Scipy (Virtanen et al., 2020), Numpy (Harris et al., 2020) and SciScripts (Malfatti, 2021b), and all plots were produced using Matplotlib (Caswell et al., 2020). Spikes were detected and clustered using SpyKING Circus (Yger and Marre, 2019), and visual inspection was performed using Phy (Rossant et al., 2016).

GPIAS signal was bandpass filtered from 70 to 400Hz for piezoelectric recordings and lowpass filtered below 50Hz for accelerometer recordings. Data was cut 200ms around the loud pulse onset. For accelerometer recordings, the absolute values of the three axes were averaged. The 9 Gap-startle trials of the same frequency band were averaged, as were the 9 Startle trials. The instantaneous amplitude of the signal was calculated as the magnitude of the analytic representation of the averaged signal using the Hilbert transform. The amplitude of the response was defined as the mean instantaneous amplitude 100ms after the loud sound pulse subtracted by the mean instantaneous amplitude 100ms before the loud pulse, which corrects for baseline offsets. The GPIAS index was calculated as

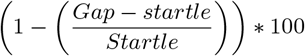

where *Startle* is the amplitude of response to Startle trials and *Gap* − *startle* is the amplitude of response to Gap-startle trials. The most affected frequency for each animal was calculated as the frequency with the greatest index shift from before to after noise exposure. Group data is shown as boxplots, where horizontal lines show the median, triangles show mean, circles show outliers and whiskers bounding 99% of the data points. Comparisons between treatments were done using two-tailed paired Student’s t-test, Bonferroni-corrected for the number of frequency bands tested. On screening GPIAS capability before including animals into the study, animals that did not show a startle suppression of at least 30% (Li et al., 2013) in Gap-startle vs Startle trials for all frequencies were re-tested on the next day only on those frequencies. Animals that still did not show a startle suppression by the silent gap of at least 30% at least two frequencies were excluded from further experiments.

ABR recordings were filtered using a 4th order butterworth digital bandpass filter (600-1500Hz), and data was sliced 3ms before to 9ms after each sound pulse onset and the 529 trials were averaged. ABR peaks were detected in the highest intensity response as values one standard deviation (SD) above the mean, larger than the previous value, and larger or equal to the next value. Next, each decreasing intensity was screened for peaks where a “valid peak” follows the above criteria and, in addition, has to be preceded by a peak in the previous intensity, displaying an increased latency compared to the peak in the higher intensity response. Hearing threshold was defined as the lowest sound intensity where a peak can be detected following the above criteria. If the threshold is defined as 35dBSPL, the animal’s actual hearing threshold was considered as ≤ 35dBSPL. As for GPIAS results, group data is shown as boxplots, where horizontal lines show the median, triangles show mean, circles show outliers and whiskers bounding 99% of the data points. Data is reported as mean ± standard error of the mean (SEM), and Student’s t-test, two-tailed, unequal variance was applied to compare pairwise differences. The reported p-values were bonferroni-corrected when the same dataset was used for multiple comparisons.

Spikes from unit recordings were detected and clustered using the following parameters: 4th order butterworth digital bandpass filter from 500 to 14250Hz; detect negative spikes; single threshold from 2 ∼ 4.5× SD; 3 features per channel. Peri-stimulus time histograms (PSTHs) were calculated by summing occurrence of spikes in a time window of 100 ms around each TTL (50ms before and 50 ms after the TTL) and presented as number of spikes per time, where each bin corresponds to 1ms. Units were classified as responding units as described by Parras et al. (2017). Spike rate was calculated as spike events per second along all the recordings (including the stimulation period). The firing rate of each unit was calculated for each frequency and intensity tested, and plotted as frequency-intensity-firing rate pseudocolor rectangular grid plots, then firing rate was bilinearly interpolated, upsampling 3x in frequency and intensity dimensions. Unit tuning width was calculated as the mean of the normalized firing rate for each frequency tested at 80dB, therefore, higher values represent broader tuning curves. Unit best frequency was defined as the sound frequency that elicited the highest firing rate. Group data is reported as mean ± SEM, and paired two-tailed Student’s t-test with unequal variance was applied to compare firing rate between neurons. Correlation between unit features (firing rate, tuning width and best frequency) was calculated as Pearson correlation coefficient and p-value for testing non-correlation.

## Results

### Inhibition of CaMKII *α*-hM4Di positive DCN cells decreases tinnitus perception

To investigate perception of noise-induced tinnitus in mice using operant tasks can be challenging and has led to the development of a modified startle suppression task for rodents such as mice and guinea pigs (Turner et al., 2006; Longenecker and Galazyuk, 2012, 2016; Longenecker et al., 2018; Park et al., 2020). Here we initially screened for capability to carry out the gap prepulse inhibition of acoustic startle (GPIAS Turner et al., 2006; Yang et al., 2007). Mice were acclimatized and habituated to the test equipment before subjected to GPIAS (Figure 1A) testing the capability of detecting a short (40ms) silence in background noise (60dBSPL) 100ms prior to a loud startle pulse (105dBSPL, 50ms duration, Figure 1D), thereby suppressing the acoustic startle reflex by at least 30% (Li et al., 2013). Six different frequency bands were pseudorandomly presented with the startle pulse (Startle session) or the silence in noise (Gap-startle session) and the startle suppression index was calculated for each frequency. Mice (P26) not showing gap-detection capabilities for at least two frequencies were excluded from further experiments (4/34 mice, 11.8%; Li et al., 2013).

Next, mice were injected bilaterally with viral vectors to transduce expression of inhibitory Designer Receptors Exclusively Activated by Designer Drugs (DREADDs; Armbruster et al., 2007) based on mutated muscarinic (M4) receptors (rAAV5/CaMKII*α*-HA-hM4D(Gi)-IRES-mCitrine, or for control experiments only a fluorescent protein, rAAV5/CaMKII*α*-eYFP) containing the CaMKII*α* promoter, in the DCN. Mice were returned to their home cage and left approximately one month (Figure 1A) for adequate hM4Di expression in CaMKII*α* expressing cells, comprising both excitatory and some inhibitory cell populations (Malfatti et al., 2021). Hearing threshold was evaluated by recording auditory brainstem responses (ABRs, Figure 1B-C) three days prior to noise exposure (1h, 90dB-SPL, 9-11kHz filtered uniform white noise, followed by 2h in silence) under anesthesia in order to induce tinnituslike behavior (Winne et al., 2020). Recording of ABRs were repeated three days after noise exposure to examine any potential hearing threshold shift (Figure 1A; Winne et al., 2020), as the aim was to study tinnitus mechanisms unrelated to persistent hearing loss. ABRs showed no significant difference in hearing threshold (hM4Di noise exposed: 41±0.9dBSPL, n = 11 mice; eYFP noise exposed: 47±1.1dBSPL; n = 7 mice, *p >* 0.08 for all frequencies tested, Figure 1C).

**Figure 1:**
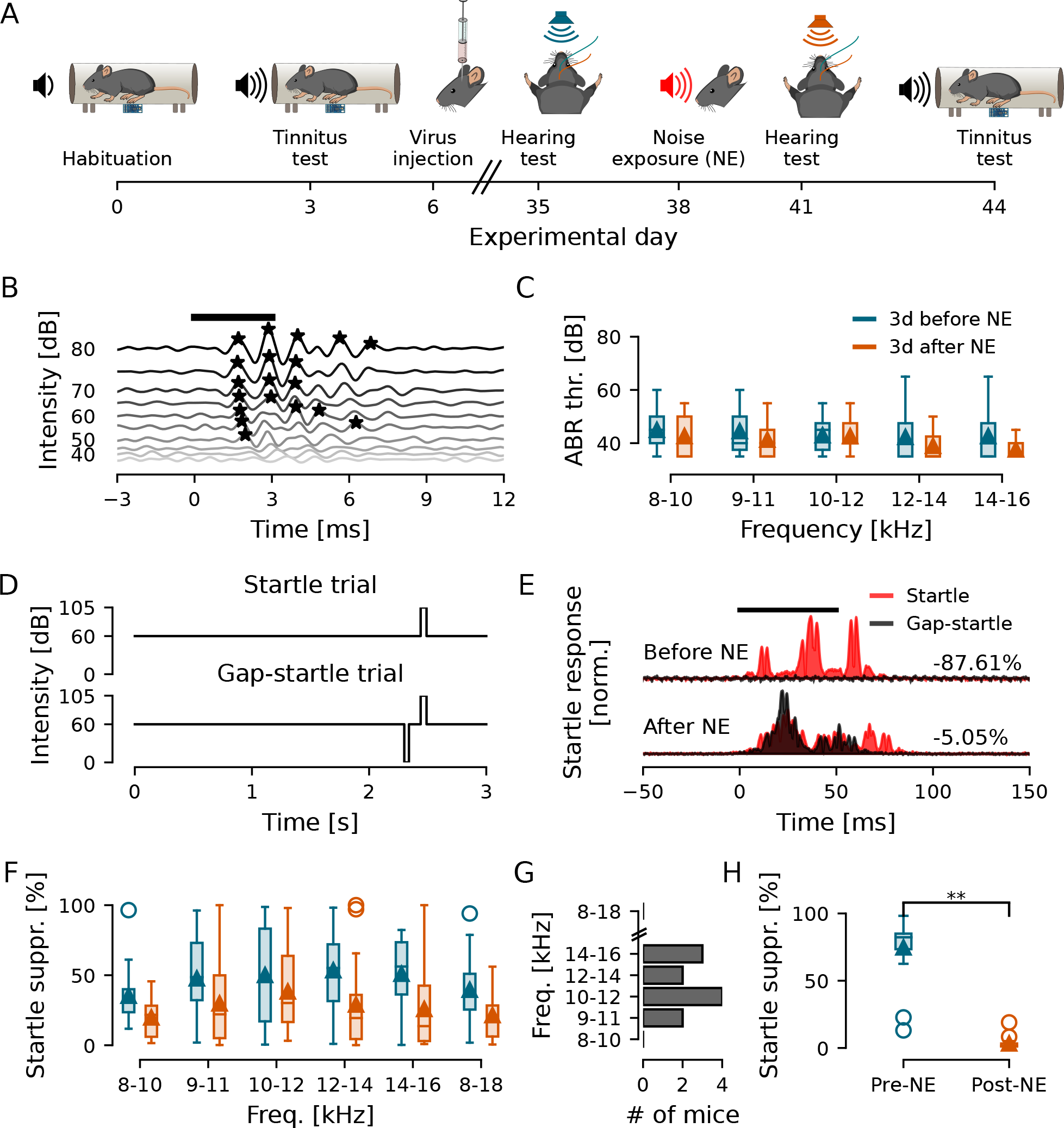
Noise exposure induces tinnitus without causing hearing loss. A) Experimental timeline. B) ABR representative example for 8-10kHz frequency presented from 80 to 35dBSPL. Response peaks are marked with black asterisks. The animal’s hearing threshold for this frequency was defined at the last intensity with an identified peak, in this example, 50dBSPL. C) Group hearing thresholds for each frequency tested before (blue) and after (orange) noise exposure (NE). D) Schematic drawn of the gap and no-gap protocols. E) Representative GPIAS recording of a mouse showing 87.6% suppression of acoustic startle before and 5.1% suppression after noise exposure when comparing no-gap (red) and gap (black) responses, indicating tinnitus-like behaviour for the tested frequency (9-11kHz). F) GPIAS group performance before (blue) and after (orange) noise exposure. G) Histogram showing the number of animals in function of the frequency with the greatest decrease in GPIAS performance. H) GPIAS group performance at the most affected frequency of each animal (n = 18 mice). **: p = 5.6e-09.

Next tinnitus-like perception was tested using GPIAS, with test rationale that if the animal has noise-induced tinnitus the animal will fail to perceive the silent gap (at a particular frequency), and thereby show lower gap-induced suppression of startle (Figure 1D-E). When measuring the GPIAS response after noise exposure, mice received an i.p. injection of NaCl (same volume as for CNO treatment, 10*µ*l/g), 30 min before the test, to perform the same procedures as for when subsequently activating inhibitory DREADDs. Group data of GPIAS indices did not reveal any particular frequency more affected by noise exposure (Figure 1F, n = 18 mice, *p>* 0.051 for all frequencies). Therefore we report the most affected frequency band, with largest change in startle suppression before and after noise exposure (Figure 1G), as parameter for tinnitus (Winne et al., 2020). Thereby our model for noise-induced tinnitus showed that loud noise exposure could induce tinnitus-like responses in mice (n = 18 mice, *p* = 5.6*e* − 09; Figure 1H) without a permanent hearing threshold shift.

Next we investigated if chemogenetically decreasing neuronal activity of the DCN can temporarily reduce tinnitus perception. For this, mice bilaterally expressing hM4Di DREADDs, or YFP for the control group, received an i.p injection of low dose CNO (0.5mg/kg) 30 min prior (Guettier et al., 2009) to the repeated GPIAS test session (Figure 2A). Mice expressing hM4Di in the DCN, that had showed tinnitus responses after noise exposure (NE - initial startle suppression: 80.2±2.3%; post NE with NaCl injection: 3.1±0.7%; n=11 mice, *p* = 9.4*e* − 09) showed a significant improvement in detecting the silent gap under the effect of CNO compared to NaCl (31.8±7.3%; *p* = 0.038; Figure 2B). Some mice showed drastic improvement as seen by the two outliers (Figure 2B), while others showed more moderate improvement (post NE with NaCl injection: 3.5±0.8; post NE with CNO injection: 21±3.2; n=9; *p* = 0.018). The control group, expressing eYFP, also with tinnitus responses after noise exposure (initial startle suppression: 67.2±12.2%; post NE with NaCl injection: 3.9±2.4%; n=7, *p* = 0.016) showed no improvement after CNO injection compared to NaCl (2.6±1%; *p* = 0.696, Figure 2C). This indicates that lowering the activity of CaMKII*α*-hM4Di positive cells in the DCN can acutely and partially ameliorate tinnitus.

**Figure 2:**
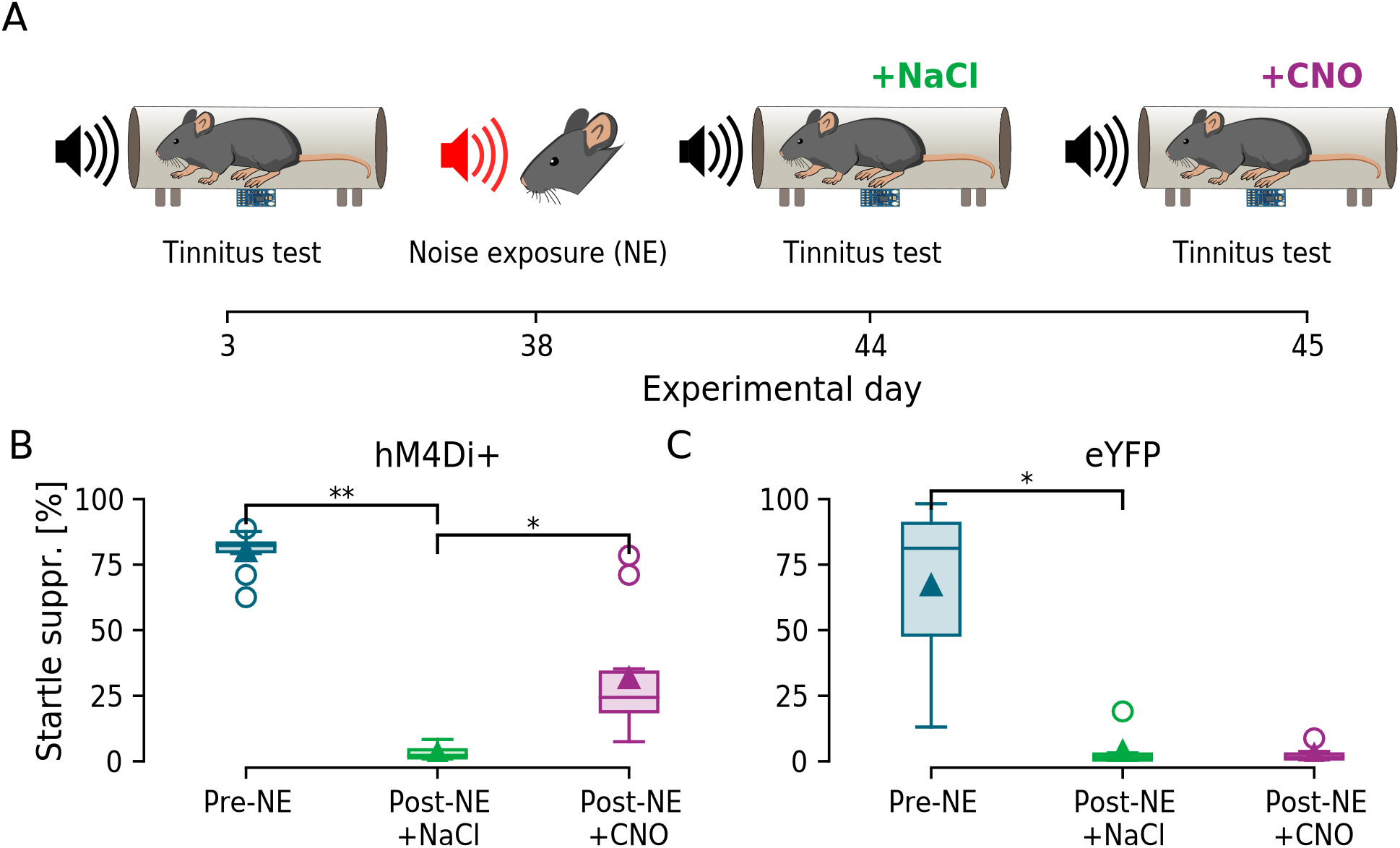
Inhibition of DCN CaMKII *α*-hM4Di positive cells after noise exposure decreases tinnitus-like behaviour. A) Schematic GPIAS recordings timeline. B) GPIAS group performance showed that animals expressing CaMKII*α*-hM4Di decreased startle suppression after noise exposure (n = 11 mice) and increased startle suppression when under the effect of CNO. C) GPIAS control group performance (expressing enhanced yellow fluorescent protein, eYFP) showing that although presenting tinnitus-like responses after noise exposure (n = 7 mice) no difference can be observed between NaCl and CNO treatments (p = 0.696). *: p *<* 0.05; **: p = 9.4e-09.

### Unit recordings confirms hM4Di expressing cells chemogenetically decrease firing

We next wanted to understand how lowering the activity of CaMKII*α*-hM4Di positive cells affected the whole DCN circuitry and therefore performed in vivo unit recordings in the presence of CNO. Recent work has shown CNO to not pass the blood-brain barrier (Gomez et al., 2017), instead reverting back to clozapine when administered (Jendryka et al., 2019) but with the ability to activate DREADD receptors at very low concentrations and avoiding off target effects (Cho et al., 2020). Therefore it was also important to directly confirm that CNO injections decreased DCN neuronal activity. DCN unit activity was recorded in response to short sound pulses (3ms; 8-10, 9-11, 10-12, 12-14 and 14-16kHz filtered uniform white noise) at different sound intensities (80-35dBSPL, 5dB-SPL decreasing steps; presented at 10Hz). Spontaneous (5min) and sound-evoked activity was recorded using a 16-channel single-shank silicon probe lowered into the left DCN (Malfatti et al., 2021) in response to auditory stimuli following NaCl and CNO i.p. injections (30 min prior to recordings, Figure 3A). A total of 224 units were isolated from 18 noise-exposed mice. Units were analyzed for firing rate and best frequency (frequency eliciting the maximum firing rate) in response to different narrow-band frequencies at different sound pressure levels (Figure 3B, see Table 1). Administration of CNO significantly decreased the average firing rate in hM4Di expressing animals in response to 80dBSPL at best frequency (NaCl: 15.85±1.95Hz vs. CNO: 8.96±1.53Hz, p = 1.3e-04, Figure 3C, left). Examining units from hM4Di+ mice in detail showed 96/122 units decreased firing rate (66±2% decrease in firing frequency; Figure 3C insets; Additional Figure 1A, middle) and 26/122 units increased firing rate following CNO administration (132±28% increase; Figure 3C insets; Additional Figure 1A, right). In control animals expressing eYFP, CNO injections did not significantly change the average firing rate of units (NaCl: 14.36±1.67Hz vs. CNO: 13.21±1.62Hz, n = 102 units from 7 mice, p = 0.4, Figure 3C, right). As auditory neurons are developmentally tuned to respond better to certain frequencies, we further analyzed tuning width and any change in best frequency of each unit. For tuning width, lower values represent narrower frequency response peaks. Here we found an average decrease in tuning width following CNO administration (0.78±0.01 to 0.74±0.01, p = 0.019, Figure 3D, left), but after closer examination 71/122 (58%) units decreased while 51/122 (42%) increased tuning width in response to the short sound pulses tested (Figure 3D insets; Additional Figure 1B). No significant changes were observed in control eYFP animals (0.67±0.01 to 0.69±0.01, p = 0.094, Figure 3D, right). Finally, we tested if units changed to what frequency they display maximum firing rate (best frequency) after CNO injection. Data showed a small but significant average increase in best frequency (12.16±0.22Hz to 12.83±0.21Hz, p = 0.026, Figure 3E, left), with 57/122 (47%) increasing, 30/122 (24%) decreasing, and 35/122 units (29%) maintaining the same best frequency for both treatments (Figure 3E insets; Additional Figure 1C). Taken together, electrophysiological data shows that inhibition of CaMKII*α*-hM4Di positive DCN cells indeed lowers the average firing rate of DCN neurons, as well as, affecting tuning width and best frequency in the DCN circuitry, which may decrease the tinnitus perception as seen by behavioral improvement of GPIAS after CNO administration.

**Table 1:** Firing rate, tuning width and best frequency features for each experimental group (NE hM4Di+ - animals exposed to noise expressing CaMKII*α*-hM4Di, n=11 mice; or NE+CNO hM4Di - animals exposed to noise under effect of CNO, expressing CaMKII*α*-hM4Di, n=6 mice) and each respective control (NE eYFP - animals exposed to noise expressing CaMKII*α*-eYFP, n=7 mice; or NE+CNO eYFP - animals exposed to noise under effect of CNO, expressing eYFP, n=6 mice) represented as mean standard error of the mean (SEM). Unit responses are further subdivided based on the applied treatment (NaCl or CNO) and on the CNO response in relation to NaCl (All - all units; Decreased and Increased - units that show a decrease or an increase in that feature under effect of CNO, respectively).

| | Firing rate (Hz; mean ± SEM) | | | | | |
| --- | --- | --- | --- | --- | --- | --- |
|  | All |  | Decreased |  | Increased |  |
|  | NaCl | CNO | NaCl | CNO | NaCl | CNO |
| NE hM4Di+ | 15.848 ± 1.948 | 8.965 ± 1.526 | 17.452 ± 2.319 | 5.566 ± 1.221 | 9.925 ± 2.916 | 21.516 ± 4.819 |
| NE eYFP | 14.365 ± 1.669 | 13.214 ± 1.621 | 20.347 ± 3.039 | 9.231 ± 2.198 | 9.614 ± 1.547 | 16.377 ± 2.253 |
| NE+CNO hM4Di+ | 9.367 ± 0.669 | 8.452 ± 0.604 | 9.902 ± 0.918 | 7.056 ± 0.688 | 8.433 ± 0.885 | 10.883 ± 1.084 |
| NE+CNO eYFP | 4.812 ± 0.682 | 4.237 ± 0.59 | 5.766 ± 1.092 | 3.232 ± 0.594 | 4.043 ± 0.808 | 5.905 ± 1.141 |

| | Tuning width (a.u.; mean ± SEM) | | | | | |
| --- | --- | --- | --- | --- | --- | --- |
|  | All |  | Decreased |  | Increased |  |
|  | NaCl | CNO | NaCl | CNO | NaCl | CNO |
| NE hM4Di+ | 0.776 ± 0.012 | 0.744 ± 0.014 | 0.822 ± 0.013 | 0.718 ± 0.02 | 0.711 ± 0.02 | 0.78 ± 0.017 |
| NE eYFP | 0.671 ± 0.012 | 0.692 ± 0.013 | 0.709 ± 0.013 | 0.624 ± 0.029 | 0.65 ± 0.017 | 0.729 ± 0.009 |
| NE+CNO hM4Di+ | 0.608 ± 0.012 | 0.63 ± 0.013 | 0.592 ± 0.022 | 0.528 ± 0.024 | 0.621 ± 0.013 | 0.694 ± 0.01 |
| NE+CNO eYFP | 0.383 ± 0.02 | 0.399 ± 0.019 | 0.423 ± 0.03 | 0.363 ± 0.028 | 0.354 ± 0.026 | 0.425 ± 0.026 |

| | Best freq. (kHz; mean ± SEM) | | | | | |
| --- | --- | --- | --- | --- | --- | --- |
|  | All |  | Decreased |  | Increased |  |
|  | NaCl | CNO | NaCl | CNO | NaCl | CNO |
| NE hM4Di+ | 12.156 ± 0.223 | 12.828 ± 0.206 | 14.4 ± 0.234 | 10.533 ± 0.253 | 10.536 ± 0.24 | 14.071 ± 0.206 |
| NE eYFP | 12.369 ± 0.18 | 12.525 ± 0.231 | 13.191 ± 0.246 | 9.957 ± 0.208 | 10.936 ± 0.166 | 14.574 ± 0.134 |
| NE+CNO hM4Di+ | 12.151 ± 0.148 | 11.698 ± 0.154 | 12.862 ± 0.197 | 10.0 ± 0.102 | 10.517 ± 0.203 | 13.217 ± 0.24 |
| NE+CNO eYFP | 10.871 ± 0.206 | 10.906 ± 0.204 | 12.739 ± 0.423 | 10.087 ± 0.212 | 10.107 ± 0.198 | 12.393 ± 0.379 |

**Figure 3:**
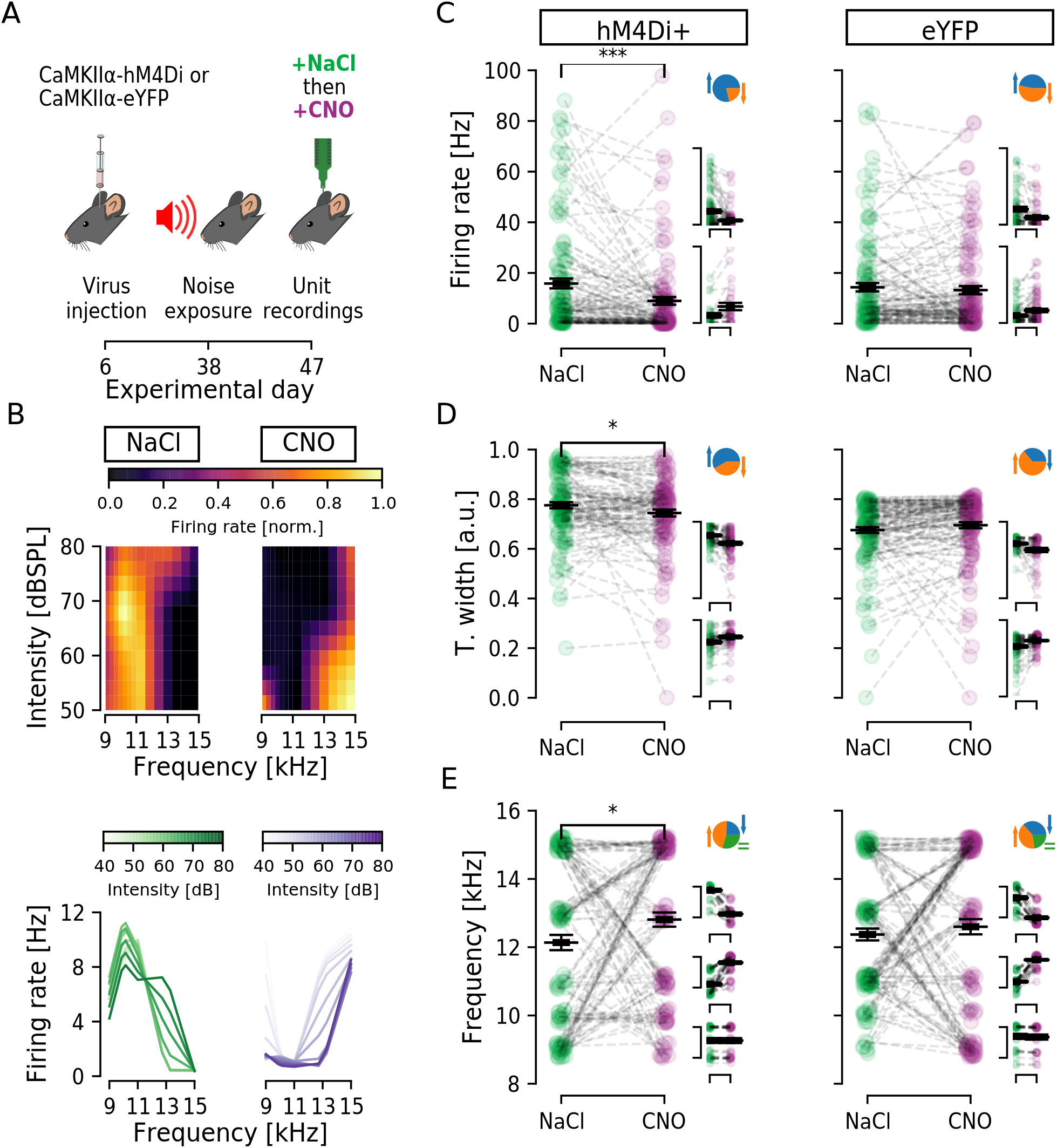
Decreasing CaMKII *α*-hM4Di positive cell activity in the DCN changes firing properties of the circuitry. A) Timeline of experiments highlighting viral injection, noise exposure and unit recordings. B) Top, firing rate (colormap) of a representative unit after NaCl (left) and CNO (right) for each intensity (lines) and each frequency (columns) tested. Bottom, a different representation of the same representative examples in the top, showing firing rate per frequency for each intensity. Data was upsampled 3 times in the intensity and frequency dimensions. C-E) Units firing rate (C), tuning width (D) and best frequency (E) for stimulation at 80dBSPL, at each unit best frequency (n = 11 mice, 122 units). Animals expressing hM4Di (left) showed a significant decrease in firing rate (C), decrease in tuning width (D) and increase in best frequency (E). Control animals expressing eYFP (right) showed no significant change in any of those parameters. Individual unit values are shown in green (NaCl) or purple (CNO) condition. Black line indicates the mean ± SEM. Insets C-E (top) shows portion of units decreasing (blue) and increasing (orange) values upon CNO administration. Inset (bottom) shows distribution of unit values divided in groups for decrease, increase or no change (for larger representation see Suppl. Figure S1). *: p *<* 0.05; ***: p = 1.3e-04.

### Decreasing CaMKII *α*-hM4Di positive DCN cells activity during noise exposure does not prevent tinnitus-like behavior

In an attempt to investigate if CaMKII*α*-hM4Di positive DCN cells are directly part of noise-induced tinnitus plasticity, we performed a new set of experiments with mice expressing CaMKII*α*-hM4Di in DCN cells now administering CNO (0.5mg/Kg, 30 min prior) during the noise exposure (1h 90dBSPL at 9-11kHz; 2h silence; Figure 4A). ABRs before and after noise exposure confirmed no indication of permanent hearing loss (n = 6 mice, *p>* 0.08 for all frequencies; Figure 4B-C) in this experimental condition. Furthermore, inhibition of CaMKII*α*-hM4Di+ DCN neurons during noise exposure did not prevent startle suppression deficit after noise exposure compared to the initial screening (n=6 mice, *p* = 5*e* − 03; Figure 4D-G), indicating that lowering CaMKII*α*-hM4Di+ DCN cell activity could not prevent noise-induced tinnitus-like behaviour. This indicates that initial noise trauma may be more related to cochlear overexcitability and that lowering activity of the DCN during loud noise does not have any overall protective effect of noise-induced tinnitus.

**Figure 4:**
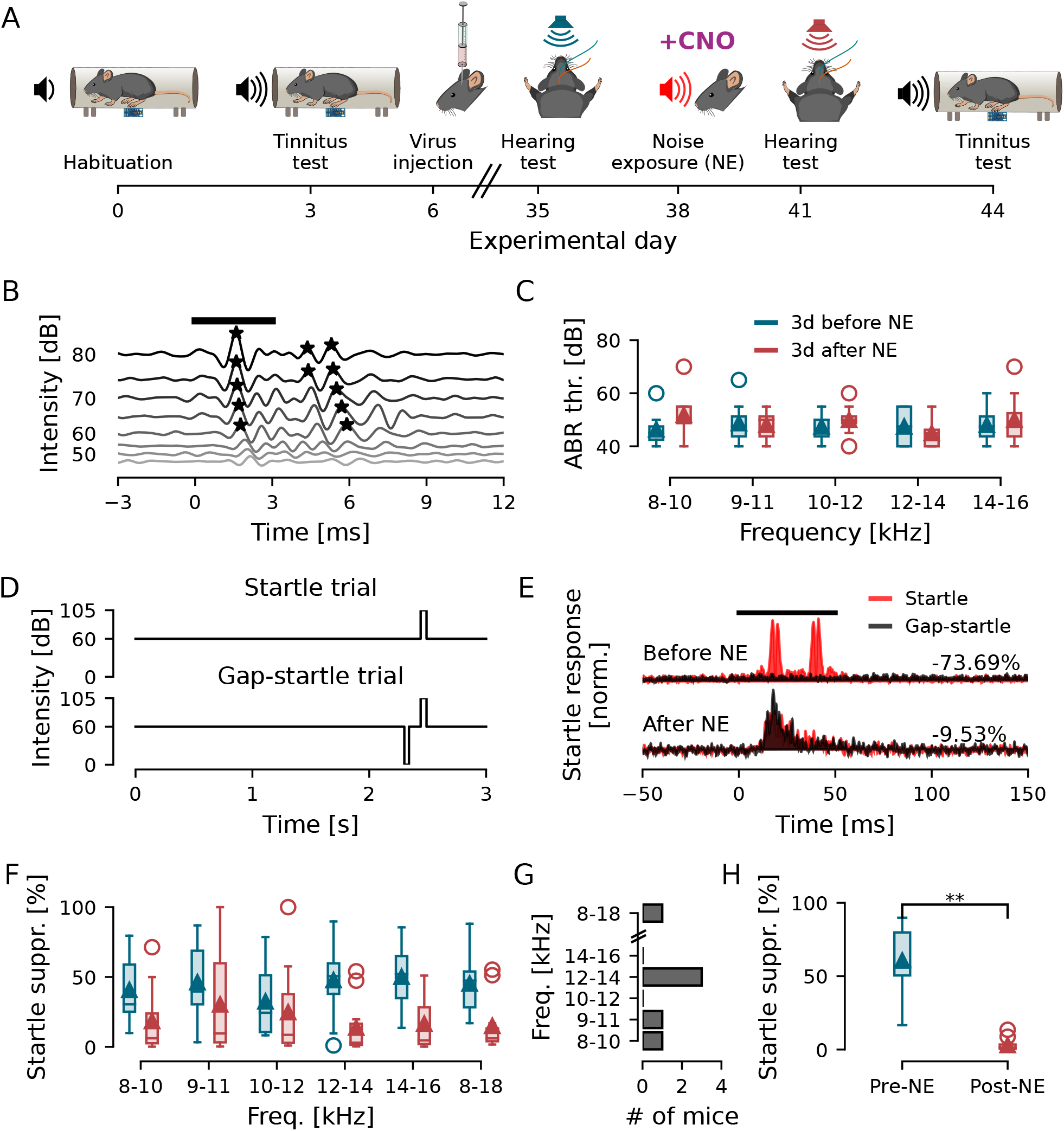
Inhibition of DCN CaMKII *α*-hM4Di positive cell activity during noise exposure does not prevent tinnitus-like behaviour. A) Timeline of experiments for hearing threshold and GPIAS recordings. B-C) Representative ABR traces and group responses for mice that received i.p. CNO injection during noise exposure. D) Schematic outline of gap and no-gap protocols. E) Representative GPIAS response. F) Group results for startle suppression of all frequencies tested before (blue) and after (dark red) noise exposure in the presence of CNO. G) Quantification of most affected frequency of each animal. H) Startle suppression of animals receiving CNO during noise exposure shows tinnitus-like behavior 12 days after noise exposure. **: P *<* 0.005

Surprisingly, in this set of experiments (Figure 5A) we found average GPIAS responses to not show any improvement in tinnitus-like responses when lowering activity of CaMKII*α*-hM4Di+ DCN cells that were inhibited during noise-exposure (hM4D1+ pre-NE: 67.5±6.8%; post-NE + NaCl: 5.3±2.2%; post-NE + CNO: 16.2±11.6%; *p* = 5.8*e* − 03 for pre-NE vs. post-NE + NaCl; *p* = 0.482 for post-NE + NaCl vs. post-NE + CNO; n = 6; Figure 5B). The control group, as expected, showed tinnitus-like responses after noise exposure (n = 6 mice, *p* = 0.023) and did not show any improvement in startle suppression after the CNO i.p. injection (eYFP pre-NE: 54.9±9.6%; post-NE + NaCl: 1.2±0.7%; post-NE + CNO: 16.5±8.8%; *p* = 0.023 for pre-NE vs. post=NE + NaCl; *p* = 0.175 for post-NE + NaCl vs. post-NE + CNO; Figure 5C). Together these experiments suggest that lowering the activity of CaMKII*α*-hM4Di positive DCN cells during noise exposure does not prevent tinnitus-like behavior, thereby CaMKII*α*+ DCN neuron activity does not appear crucial during noise exposure for triggering tinnitus. However, if CaMKII*α*+ DCN neurons were inhibited during noise exposure, the lowering of their activity using CNO in animals presenting noise-induced tinnitus no longer leads to the amelioration of tinnitus from CNO administration during the GPIAS test (Figures 1-3).

**Figure 5:**
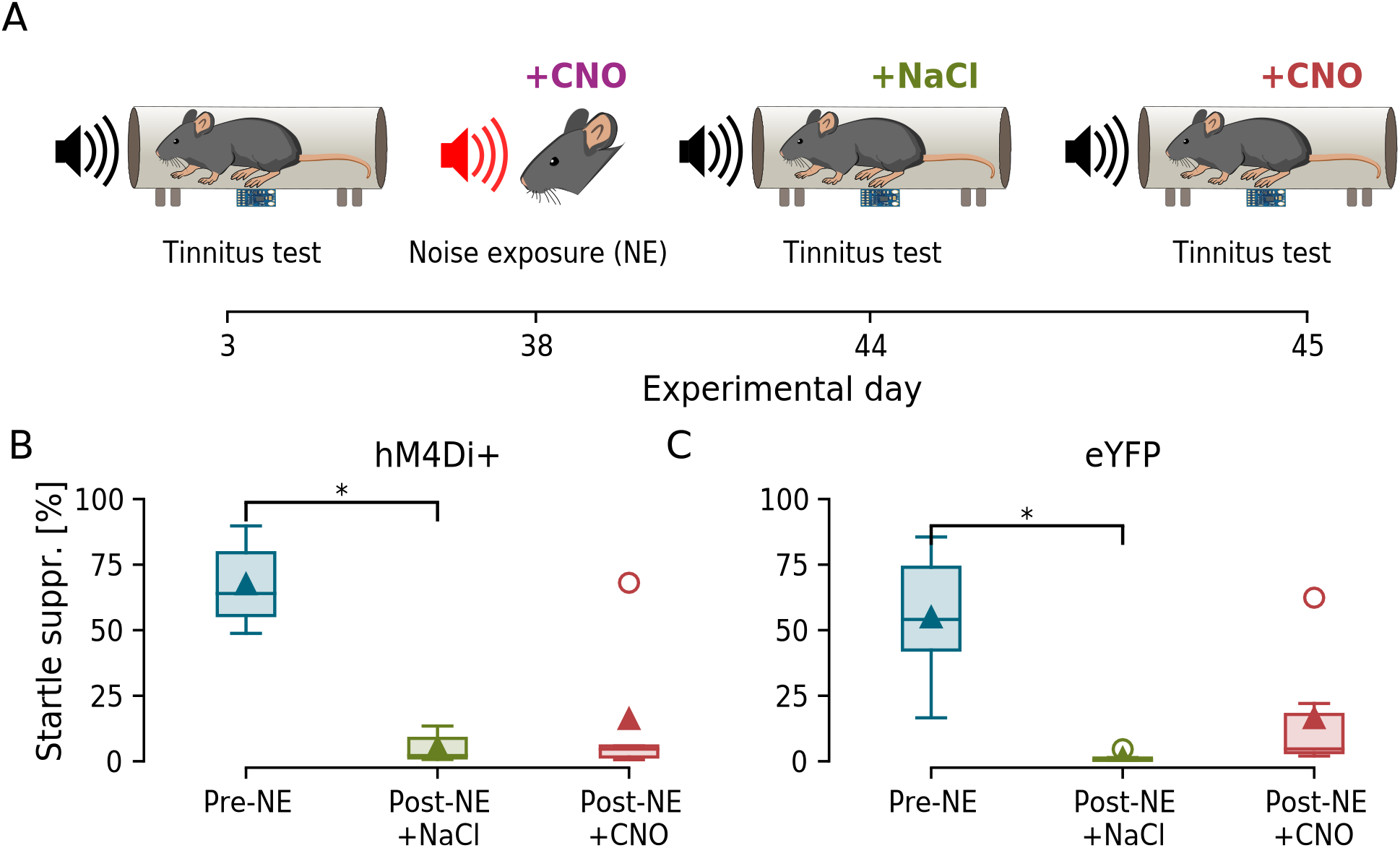
Decreasing CaMKII *α*-hM4Di positive DCN cells activity during noise exposure abolishes hM4Di-dependent recovery. A) Schematic timeline of GPIAS recordings. B) Inhibition of CaMKII*α*-hM4Di positive DCN cells during noise exposure did not prevent a decrease in the startle suppression value, indicating tinnitus (n = 6 mice), and also CNO injection during GPIAS recording after noise exposure did not recover mice startle supression (p = 0.482). C) The control group (mice expressing eYFP) showed tinnitus-like behaviour after noise exposure (n = 6 mice) and did not recover the startle supression after CNO injection (p = 0.175). *: p *<* 0.05.

### Lowered neuronal activity during noise exposure still renders units affected by CNO

To better understand the role of neural activity of the DCN at different time points of noise exposure we investigated if CNO administration lowered CaMKII*α*-hM4Di positive DCN unit activity in animals that also received CNO *during* the noise exposure (Figure 6A). Again we compared firing frequency, tuning width and best frequency in the presence of NaCl or CNO (Figure 6B, Table 1). We found that a CNO i.p. injection led to a significant decrease in firing rate (12.5±1.1Hz to 10.7±0.9Hz; n = 85 units from 6 mice; p = 4.6e-02; Figure 6C, left) in animals expressing hM4Di, but not in control animals (4.8±0.7Hz to 4.2±0.6Hz; n = 91 units from 6 mice; p = 0.195; Figure 6C, right). Also, average unit tuning width increased (0.548±0.01 to 0.587±0.01; p = 1.09e-02; Figure 6D left) and average best frequency decreased (12.6±0.2Hz to 11.8±0.2Hz; p = 4.9e-02; Figure 6E left), while the control group, expressing only eYFP, showed no significant changes in either of the parameters (p = 0.104 and 0.113, respectively; Figures 6D and E right, Table 1).

**Figure 6:**
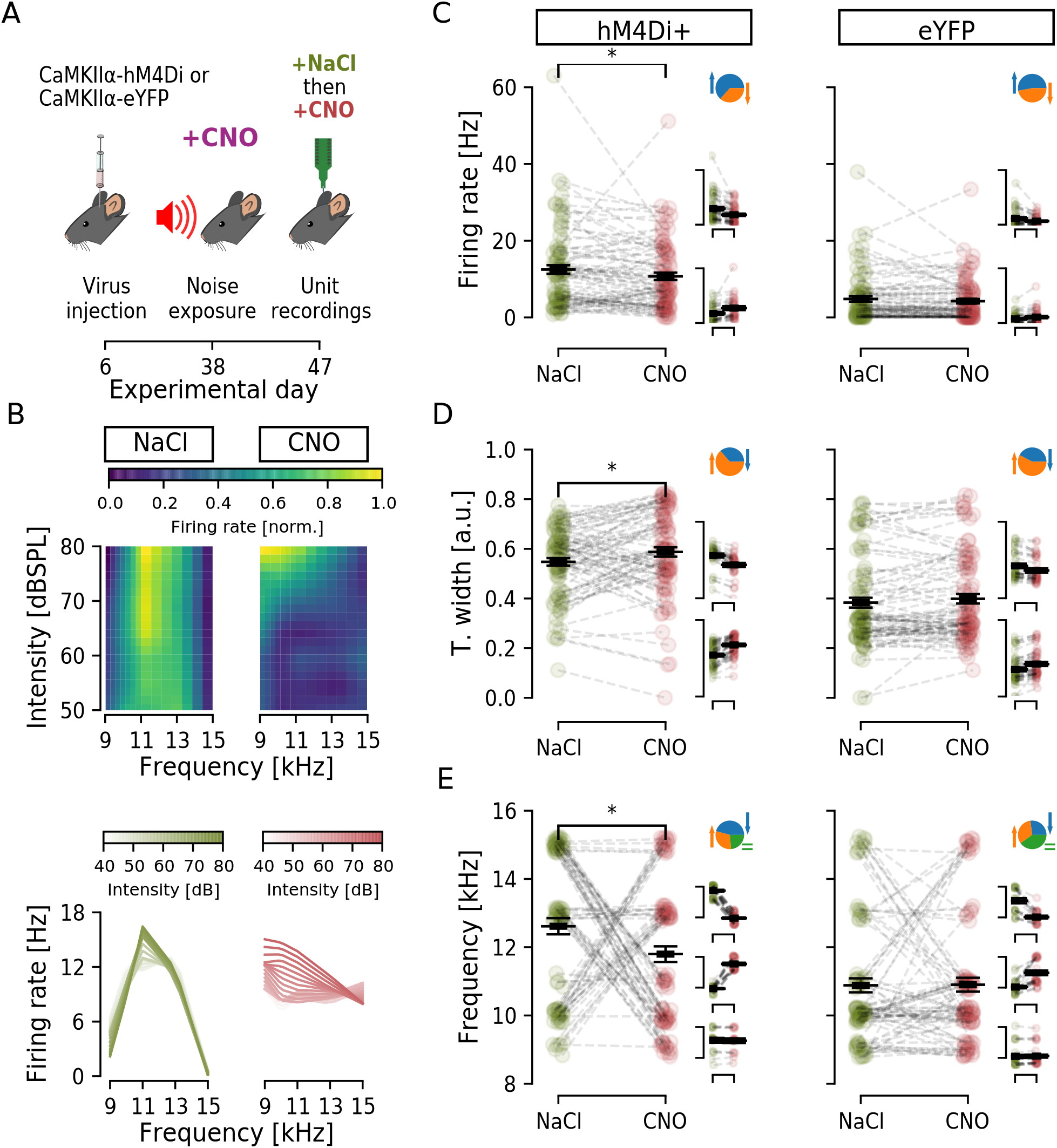
Decreasing activity of CaMKII *α*-hM4Di positive DCN cells that were also inhibited during noise exposure changes firing properties of the circuitry. A) Timeline of experiments highlighting time of viral injection, noise exposure with CNO i.p. injection and unit recordings. B) Top, firing rate (colormap) of a representative unit after NaCl (left) and CNO (right) injection for each intensity (lines) and each frequency (columns) tested. Bottom, a different representation of the same representative examples in the top, showing firing rate per frequency for each intensity. C-E) Units firing rate (C), tuning width (D) and best frequency (E) for stimulation at 80dBSPL, at each unit best frequency. Left, 85 units from mice expressing hM4Di (n = 6 mice), showing significant difference after CNO application for Firing rate, Tuning width and Best frequency. Individual unit values are shown in green (NaCl) or red (CNO) condition. Black line indicates the mean SEM. Right column, 91 units from mice expressing control eYFP (n = 6 mice), no significant difference. Insets show proportion of units decreasing (blue) or increasing (orange) parameters of each graph (see Suppl. Figure S2 for greater detail). *: p *<* 0.05.

Although the average response showed a significant decrease in firing frequency upon CNO administration, the modulation appeared bidirectional with 54 unit decreasing and 31 units increasing firing rate (Figure 6C insets; Suppl. Figure 2A). Similar results were seen for tuning width (31 units decreasing and 54 units increasing, Figure 6D insets) and best frequency (39 units decreasing, 46 units increasing, Figure 6E insets, Additional Figure 2B-C). Interestingly, the unit firing rate from animals pre-treated with CNO during noise exposure was mostly below 40kHz in these experiments, indicating a lower sample of high frequency firing units in these animals, or that typical fast spiking units fired at a lower frequency. Also, more importantly, these results show that the lack of tinnitus-like behavior improvement in the group pre-treated with CNO during noise exposure was not due to a lack of hM4Di+ neurons having an effect of decreasing activity within the DCN.

### DCN units are differently modulated by DREADDs if activity was also lowered during the noise-exposure

To better understand alterations in firing patterns of the DCN when activating inhibitory DREADDs we did correlation analysis of all obtained data. Unit recordings describe units responding to sound when CaMKII*α*+ neurons of the DCN circuit had the firing frequency in response to sound chemogenetically lowered, and does thus not reflect recordings from CaMKII*α*+ units only (Malfatti et al., 2021). Although CNO (0.5mg/kg) administration consistently lowered the average firing rate in animals expressing hM4Di DREADDs in DCN CaMKII*α*+ neurons, the bidirectional modulation seen when looking at individual DCN unit responses to sound after CNO administration made us specifically question whether any correlation exist between firing rate, tuning width and best frequency in response to CNO (Table 2). Here we display the units features as 3-dimensional plots for hM4Di+ and control animals (Figure 7) that received CNO during GPIAS to ameliorate from tinnitus (Figure 7B) and from hM4Di+ and control animals receiving CNO both during the noise-exposure and during GPIAS (Figure 7C) and examined any correlation between unit parameters using Pearson correlation coefficient (r), with the p-value testing non-correlation (Table 2). We divided our analysis in comparing all units for each experimental group, but also subdivided analysis in units decreasing or increasing firing upon CNO administration, respectively (Table 2). In the first set up experiments, trying to recover tinnitus-like behaviour (Figures 1-3), we found no correlation between average firing rate and best frequency for either experimental group, suggesting that decreasing CaMKII*α*-hM4Di+ cells firing rate does not alter units tuning to a certain frequency. Firing rate and tuning width appeared equally correlated in the presence of NaCl or CNO, indicating that lowering CaMKII*α*-hM4Di+ cells activity using DREADDs does not decouple the existing correlation between firing rate and tuning width. However, when splitting data into units either decreasing (96/122) or increasing (26/122) firing rate in response to CNO it appears that units decreasing firing rate upon CNO administration no longer correlate with tuning width, meaning that units showing low firing rate do not necessarily have a low tuning width (Table 2; Additional Figure 1A-B). In experiments where CNO was given during the noise exposure in an attempt to prevent tinnitus-like behaviour (Figure 4-6), we instead noted that firing rate was not correlated with tuning width. Interestingly, CNO administration during unit recordings appeared to recover this missing correlation (Table 2). This could indicate that CNO during noise-exposure can influence lateral inhibition within the DCN circuitry, since the firing rate is no longer coupled to the tuning of response to sound, for example units responding with a low firing rate but broadly to neighboring frequencies.

**Figure 7:**
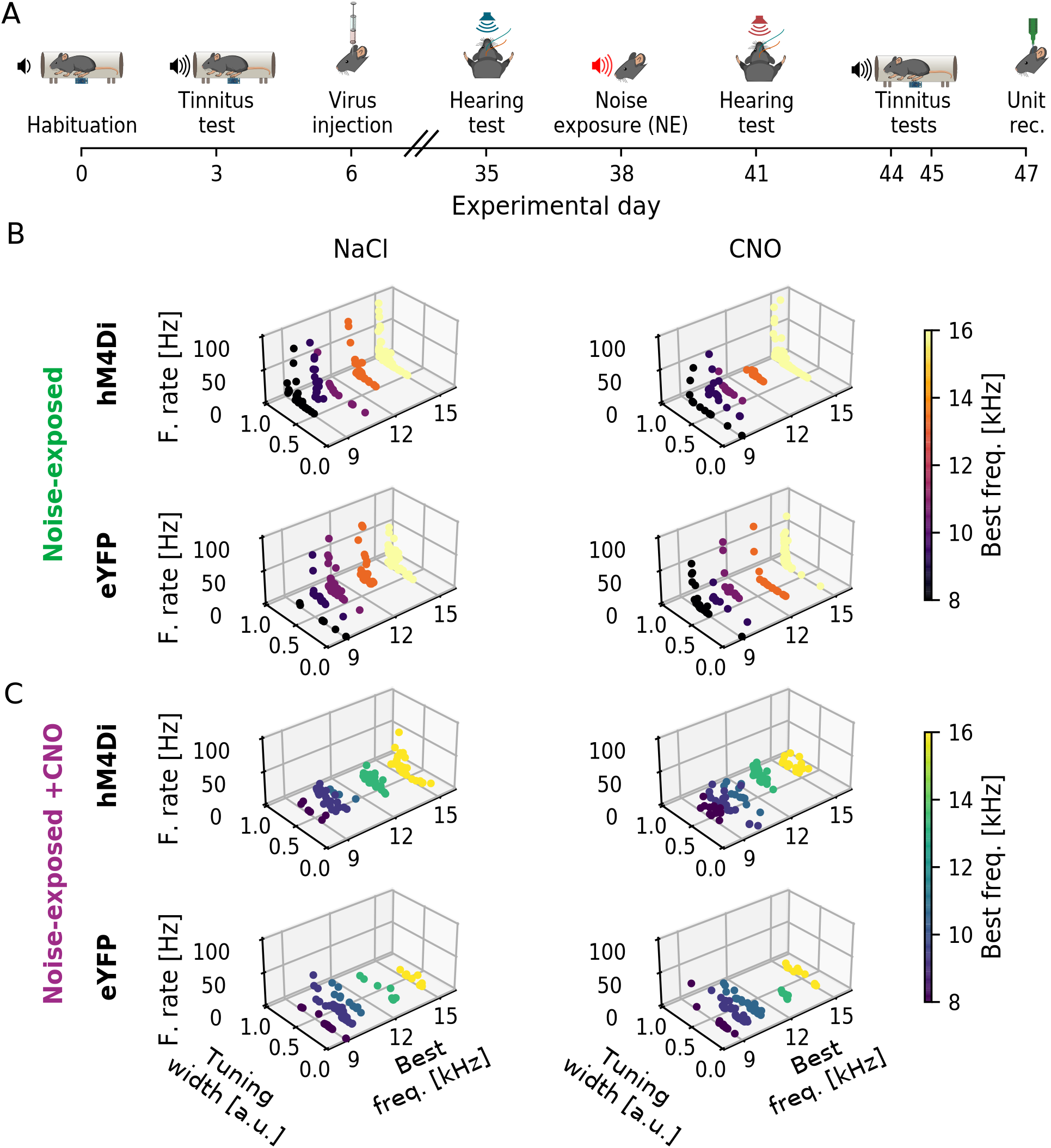
Three-dimensional scatter plots of Firing rate, Tuning width and Best frequency of DCN units of noise exposed hM4Di+ or eYFP+ animals in the presence of NaCl or CNO. A) Full experimental timeline. B) 3D scatters representing each unit by *firing rate x tuning width x best frequency* for hM4Di (experimental; top) and eYFP (control; bottom) animals under NaCl (left) or CNO (right) treatment. C) Same as B for experiments where animals were administered CNO (0.5mg/kg) 30 minutes prior to unit recordings. Colors represent the best frequency response between 8-16kHz.

**Table 2:** Correlation pairs of firing rate (FR), tuning width and best frequency features for each experimental group (NE hM4Di+ - animals exposed to noise expressing CaMKII*α*-hM4Di, n=11 mice; or NE+CNO hM4Di - animals exposed to noise under effect of CNO, expressing CaMKII*α*-hM4Di, n=6 mice) and each respective control (NE eYFP - animals exposed to noise expressing CaMKII*α*-eYFP, n=7 mice; or NE+CNO eYFP - animals exposed to noise under effect of CNO, expressing eYFP, n=6 mice) represented as Pearson correlation coefficient (r) and p-value for testing non-correlation (p). Unit responses are further subdivided based on the applied treatment (NaCl or CNO) and on the firing rate change under CNO in relation to NaCl treatment (All - all units; Decreased and Increased - units that show a decrease or an increase in firing rate under effect of CNO, respectively).

| Firing rate x Best freq. (r, p) |  |  |  |  |  |  |
| --- | --- | --- | --- | --- | --- | --- |
|  | All |  | Decreased FR after CNO |  | Increased FR after CNO |  |
|  | NaCl | CNO | NaCl | CNO | NaCl | CNO |
| NE hM4Di+ | 0.064 1.000 | 0.059 1.000 | 0.085 1.000 | -0.094 1.000 | -0.031 1.000 | 0.047 1.000 |
| NE eYFP | 0.164 0.639 | 0.142 1.000 | 0.119 1.000 | 0.37 0.053 | 0.067 1.000 | -0.081 1.000 |
| NE+CNO hM4Di+ | -0.081 1.000 | -0.024 1.000 | 0.022 1.000 | -0.061 1.000 | -0.323 0.058 | 0.007 1.000 |
| NE+CNO eYFP | 0.069 1.000 | 0.023 1.000 | 0.003 1.000 | 0.229 1.000 | 0.054 1.000 | -0.224 1.000 |

| Firing rate x Tuning width (r, p) |  |  |  |  |  |  |
| --- | --- | --- | --- | --- | --- | --- |
|  | All |  | Decreased FR after CNO |  | Increased FR after CNO |  |
|  | NaCl | CNO | NaCl | CNO | NaCl | CNO |
| NE hM4Di+ | 0.479 2.2e-07* | 0.352 6.1e-04* | 0.504 1.5e-06* | 0.264 0.083 | 0.413 0.324 | 0.484 0.108 |
| NE eYFP | 0.378 1.6e-04* | 0.445 2.5e-06* | 0.557 1.1e-04* | 0.47 3.0e-03* | 0.349 3.2e-02* | 0.429 2.3e-03* |
| NE+CNO hM4Di+ | 0.045 1.000 | 0.246 5.3e-03* | 0.099 1.000 | 0.293 9.9e-03* | -0.109 1.000 | 0.14 1.000 |
| NE+CNO eYFP | 0.47 5.0e-05* | 0.481 2.9e-05* | 0.469 1.1e-02* | 0.39 0.074 | 0.492 2.4e-02* | 0.596 1.4e-03* |

| Tuning width X Best freq. (r, p) |  |  |  |  |  |  |
| --- | --- | --- | --- | --- | --- | --- |
|  | All |  | Decreased FR after CNO |  | Increased FR after CNO |  |
|  | NaCl | CNO | NaCl | CNO | NaCl | CNO |
| NE hM4Di+ | 0.301 6.7e-03* | 0.352 6.3e-04* | 0.201 0.45 | 0.314 1.6e-02* | 0.591 1.4e-02* | 0.249 1.000 |
| NE eYFP | 0.198 0.252 | 0.252 4.7e-02* | 0.182 1.000 | 0.138 1.000 | 0.247 0.378 | 0.263 0.27 |
| NE+CNO hM4Di+ | 0.069 1.000 | 0.137 0.522 | 0.064 1.000 | 0.108 1.000 | 0.086 1.000 | 0.181 1.000 |
| NE+CNO eYFP | 0.068 1.000 | 0.231 0.306 | -0.11 1.000 | 0.365 0.126 | 0.223 1.000 | 0.036 1.000 |

Interestingly CNO administration prior to noise-exposure also showed a particular loss of correlation between firing rate and tuning width in control animals, for units decreasing firing rate following CNO administration compared to NaCl (NaCl r,p: 0.469, 1.1e-02; CNO r,p: 0.39, 0.074). This suggests that CNO, converted to clozapine, could have small electrophysiological effects on the DCN circuitry that is not seen behaviorally nor in averaged data (Figure 6, Table 2). When investigating correlations between Tuning width and Best frequency we only observed correlations between the parameters in the groups with noise-exposure without pharmacological manipulation. The correlation between tuning width and best frequency was seen for units decreasing firing rate upon CNO administration, but for units that increased firing frequency upon CNO administration this correlation was lost (NaCl r,p: 0.591, 1.4e-02; CNO r,p: 0.249, 1.0). We again observed a correlation between tuning width and best frequency in control animals only appearing following CNO administration. This correlation was however lost when units were divided into increasing or decreasing firing frequency following CNO administration. Still, it highlights the possibility that clozapine has small electrophysiological effects despite the very low dose CNO used in this study, and that despite group data not being significantly different for control animals, there may be small membrane effects through binding of clozapine to certain receptors - effects that are not seen when clozapine’s main effect is activating DREADDs. Finally, we did not record from units of either experimental group (noise-exposed or noise-exposed + CNO) at any particular depth or layer, as we did not want to bias data to any particular frequency region of the DCN (Figure 8).

**Figure 8:**
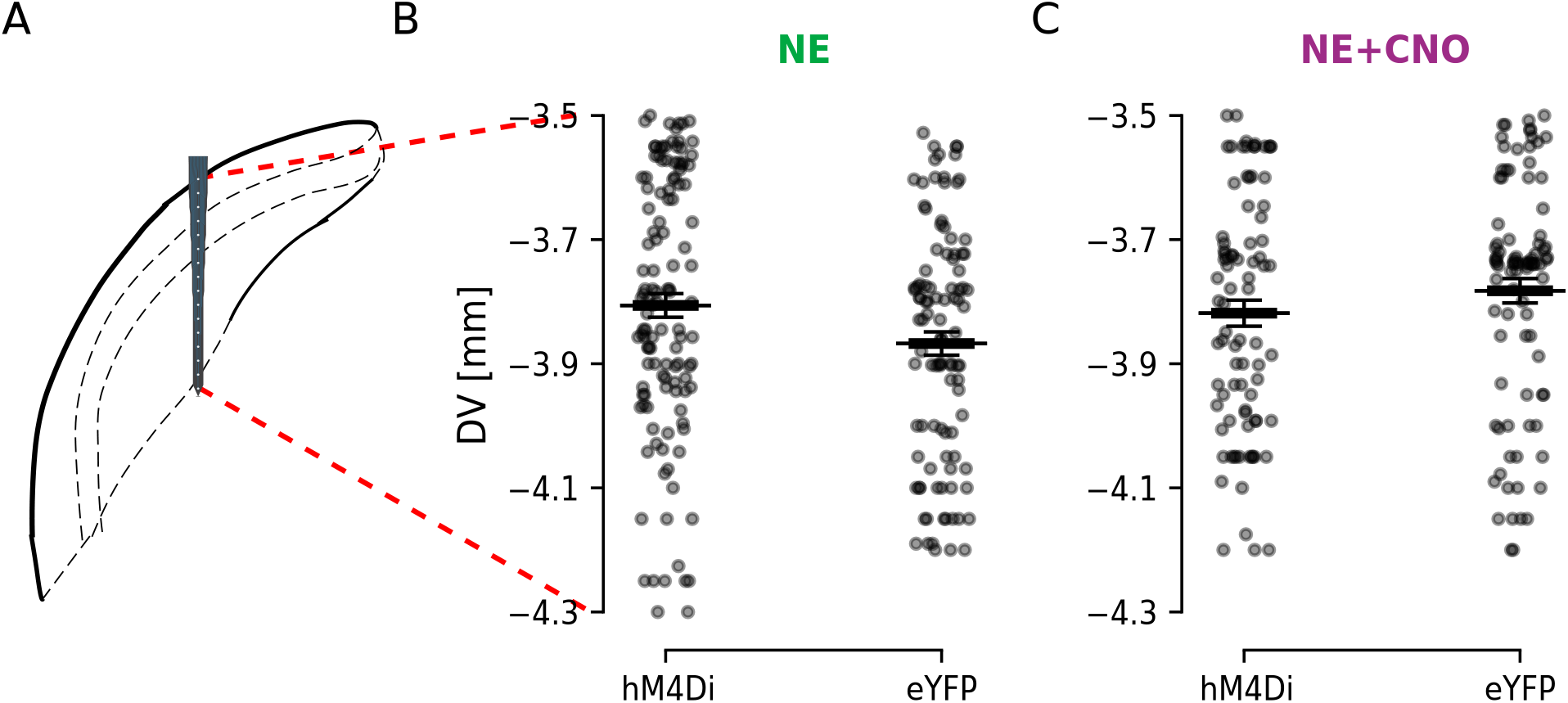
DCN unit depth profile. A) Schematic representation of the probe location within the DCN according to coordinates used highlighting the dorsoventral depth of unit recordings. B) Distribution of recorded DCN units along the dorsoventral axis for noise exposed animals expressing CaMKII*α*-hM4Di or CaMKII*α*-eYFP. C) The same as B but for experimental and control animals that were pre-treated with CNO 30 minutes before noise exposure. Black bars indicate mean *±* SEM.

## Discussion

Here we found that decreasing activity of CaMKII*α*-hM4Di positive DCN cells after noise exposure can decrease tinnitus-like responses and showed that activity of the DCN is involved in the maintenance of tinnitus perception. Still, many higher areas are most likely also involved in tinnitus but this study shows that lowering neuronal activity of the dorsal cochlear nucleus might be able to counteract increased activity or gain observed in higher auditory areas after noise exposure (Asokan et al., 2018; Shore and Wu, 2019). Moreover, this DCN subpopulation does not appear to have an important role in initial tinnitus generating mechanisms, since inhibiting CaMKII*α*-hM4Di positive DCN cells during noise exposure did not prevent development of tinnitus-like response. Instead, decreasing firing capability of CaMKII*α*+ DCN neurons during noise exposure abolished CNO-dependent recovery after noise exposure. This suggests that CaMKII*α* positive neurons play an indirect role in tinnitus maladaptive plasticity but not in the temporal window of the noise exposure.

In this study we aimed to not confound mechanisms of increased neuronal activity due to noise exposure with plasticity related to partial hearing loss as several studies in children, adolescence and adults show the prevalence of noise-induced tinnitus with normal audiograms (Sanchez, 2014; Nagaraj et al., 2019; Joo et al., 2020) This is an important group to consider, especially since noise-induced tinnitus is becoming more common in youth (Mahboubi et al., 2013; Gilles et al., 2014; Nemholt et al., 2019). In comparison to other animal models of noise-induced tinnitus, our noise exposure is on the lower edge of parameters often used (Bauer and Brozoski, 2001; Heffner and Harrington, 2002; Basta and Ernest, 2004; Kujawa and Liberman, 2009; Wu et al., 2016; Yang et al., 2016; Heeringa et al., 2018; Han et al., 2019; van Zwieten et al., 2019). We used 90dBSPL for 1h followed by 2h of silence, as we have previously shown to be able to generate tinnitus-like behavior without permanent threshold shifts (Winne et al., 2020). Here we show again that noise exposure at 9-11kHz frequency does not caused tinnitus-like behavior at a particular frequency, similarly to studies of mice and guinea pigs (Coomber et al., 2014; Longenecker and Galazyuk, 2016), as well as for patients reporting noise exposure as tinnitus etiology (Zagáolski and Strek, 2014).

It is well known that the DCN circuitry presents altered firing following noise exposure (Shore and Wu, 2019). DCN cells, specially fusiform cells, can increase spontaneous activity (Baizer et al., 2012), bursting activity and synchrony (Wu et al., 2016). Still, it is not established if the DCN undergoes plasticity during a loud noise exposure. We tried to partially address this question using chemogenetics, still acknowledging that clozapine-N-oxide administration is not temporally precise. For example, it has been shown that CNO has a half-life of 2h in mice, and with biological effects lasting 6-10h (Guettier et al., 2009). Here we saw that decreasing the activity of CaMKII*α*-hM4Di positive DCN subpopulation, during the loud noise exposure, could not counteract activity of the auditory system enough to prevent tinnitus in mice. How long the CNO effects persist and potential downstream targets were not assessed in this study, and additional studies with repeated CNO administration over longer periods following tinnitus induction would be interesting to evaluate.

One interesting indirect finding of our second set of experiments was that, if CaMKII*α*-hM4Di positive DCN cells still have a role in tinnitus triggering, they are not the only subpopulation involved, since inhibiting them was not enough to prevent tinnitus. Here, mice still develop tinnitus-like responses in startle suppression tests, but since the CaMKII*α*-hM4Di positive DCN cells were inhibited during noise exposure, we can speculate that no plasticity took place in those cells, and they would not contribute to abnormal signaling in the DCN. Thereby, inhibiting those cells later in the GPIAS test did not improve the tinnitus perception. More so, several different subpopulations of DCN neurons could explain the fact that mice that recovered after CNO injection in the first set of experiments presented only a partial recovery (Figure 2B, the startle suppression was not restored to pre-noise exposure values). Decreasing activity of a subgroup of DCN neurons could still significantly improve acoustic startle suppression compared to post-noise exposure. This points to the importance of restoring inhibition in tinnitus (Richardson et al., 2012), where chemogenetic inhibition thus has the potential to alleviate tinnitus.

Recent studies have shown that clozapine-N-oxide cannot cross the blood-brain barrier, instead CNO may be reverted to the antipsychotic compound clozapine that crosses the blood-brain barrier and can bind to a variety of neurotransmitter receptors (Gomez et al., 2017). Still, Manvich et al. (2018) showed that the amount of CNO necessary to cause unspecified behavioral changes in mice or rats was 5mg/kg, which is 10x greater than the dose administered in our study. We observed no changes in GPIAS responses or unit recordings of animals injected with CNO without expressing the hM4Di receptor. Instead we saw effects specific for inhibiting CaMKII*α*-hM4Di positive DCN cells, with CNO causing a significant decrease in DCN units firing rate. Still, some units showed an increase in firing rate. Since hM4Di activity through second messengers leads to membrane hyperpolarization (Rogan and Roth, 2011), units showing an increase in firing rate after CNO injection are most likely being disinhibited (due to some of the targeted CaMKII*α*-hM4Di positive DCN cells being inhibitory, Malfatti et al., 2021). Still, the effects of CNO on best frequency was not binary (increased or decreased best frequency), as some units did not change best frequency. Interestingly we found that, in noise-exposed animals, firing rate was not correlated with best frequency, regardless if CaMKII*α*-hM4Di positive DCN cells were inhibited or not. This is probably due to the analysis of the provided stimulus, as many DCN cells may have best frequencies higher than 16kHz, and also respond differently to pure-tones (Godfrey et al., 1975; Nelken and Young, 1994). Hence, some units could have been erroneously classified, for example, as a unit with a low firing rate and broad tuning width, confusing results relating to the effect of CNO. Thereby no inferences about best frequency and tinnitus-like responses, or the tonotopicity of the DCN, have been made in this study.

Nevertheless, a neuron’s tuning width is an important feature that allows refinement of circuitry responses to stimuli. The literature is still ambiguous on how firing rate and tuning width are correlated in DCN units, since generalized inhibition by intracerebral injection of muscimol (GABAA receptor agonist) was shown to not affect tuning width in DCN units of anesthetized rats (Paolini et al., 1998); while another study showed that bicuculline (a GABA receptor antagonist) increased tuning width while GABA and muscimol decreased tuning width (Yajima and Hayashi, 1990). We found that firing rate was correlated with tuning width, except for animals where CaMKII*α*-hM4Di positive DCN cells were inhibited during noise exposure. We speculate that forcing CaMKII*α*-hM4Di+ DCN cells to keep a low firing during the noise exposure decoupled firing frequency from tuning width, and thereby only a portion of DCN cells would show hyperactivity due to noise exposure (Pilati et al., 2012; Wu et al., 2016). Interestingly, inhibiting these cells during unit recordings restored the correlation between firing rate and tuning width, perhaps by decreasing activity of cells not affected by the noise exposure, and thereby emphasizing regular activity of the remaining circuitry.

In summary, our results illustrate the complexity of the DCN circuitry and indicate that decreasing CaMKII*α*-hM4Di positive DCN cell activity may drastically change DCN circuit electrophysiology. Such changes may underlie improvements related to for example loudness of tinnitus (Kaltenbach, 2006) which is of clinical relevance as one neurological treatment effect in tinnitus patients is a decreased loudness and/or decrease of Annoyance index (Lefaucheur et al., 2017, 2020). In conclusion, our results show that CaMKII*α*-hM4Di positive DCN cells play a significant role in maintaining noise-induced tinnitus in mice, and provide a step towards better understanding the neuronal correlates of noise-induced tinnitus in patients with normal hearing threshold.

## Author Contributions

TM and BC performed the experiments; TM and MH analyzed the data; TM, BC and KEL wrote the manuscript with input from RNL.

## Acknowledgments

We would like to thank Dr. Helton Maia, Dr. George Nascimento and Dr. João Bacelo for technical advice.

## Data Availability Statement

The datasets generated and/or analyzed in the current study are available on request.

