## Supplementary Figure 1-2 for "Chemogenetically decreasing activity of the dorsal cochlear nucleus can ameliorate noise-induced tinnitus in mice"

### Supplementary figures

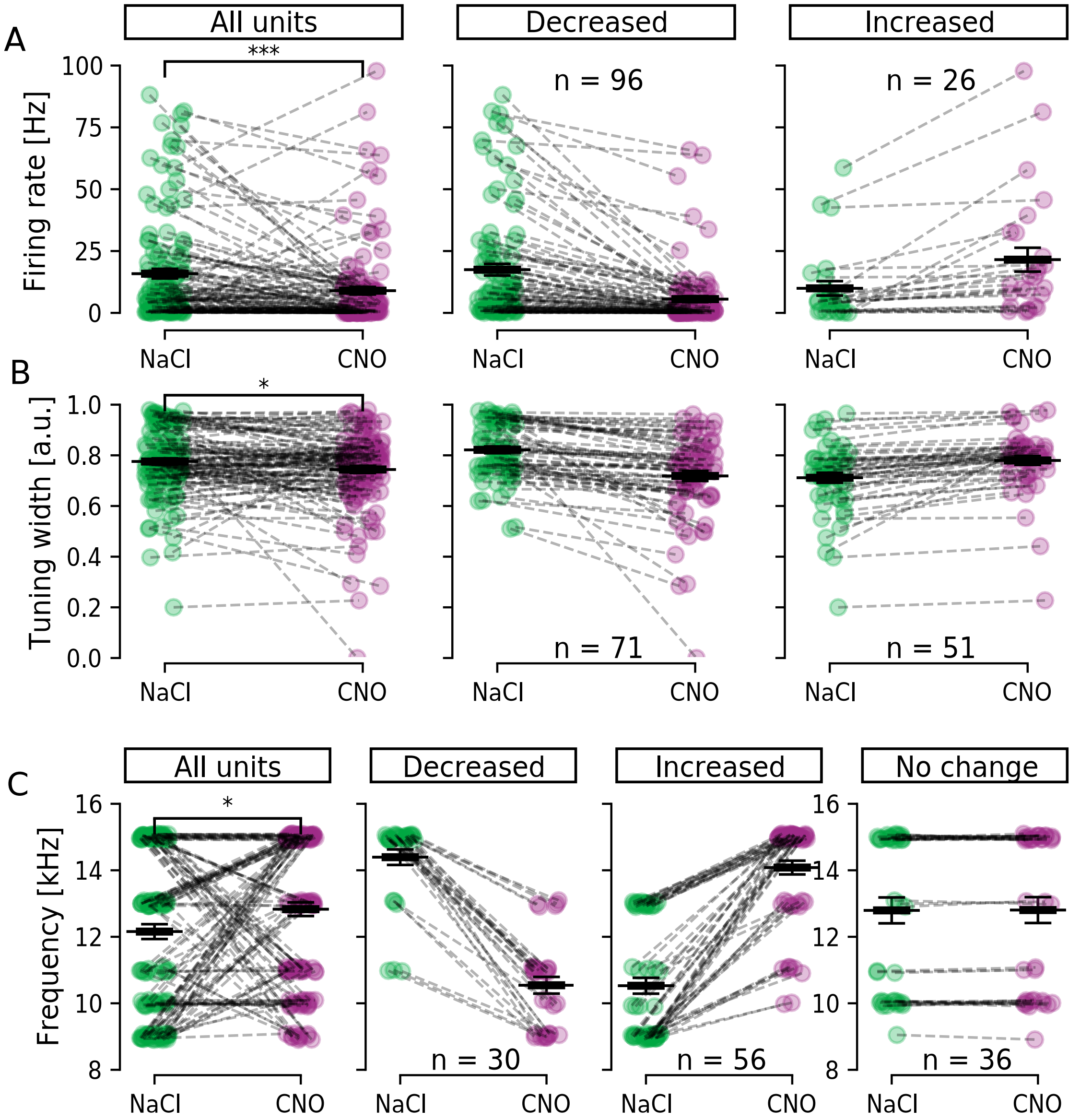


**Supplementary figure S1. Bimodal unit responses seen upon CNO administration in hM4Di+ mice**. A) Left; Firing rate in response to 80dBSPL at best frequency for all units (n = 122) from hM4Di+ mice in response to NaCl or CNO. Middle; Only units decreasing (n = 96) firing rate upon CNO administration. Right; Units increasing (n = 26) firing rate after CNO administration. B) Same as 'A' but for Tuning width, with units decreasing (n = 71) and increasing (n = 51) tuning with after CNO administration. C) Same as 'A' for representation of Best frequency in kHz, with units decreasing (n = 30), increasing (n = 56) or maintaining (n = 36) Best frequency response upon CNO administration. Note that units responding to sound do not need to be CaMKII$\alpha$+, the unit altered firing properties are in response to sound when CNO is decreasing activity of CaMKII$\alpha$+ units of the DCN circuit. *: $p < 0.05$; ***: $p = 1.3e\text{-}04$.

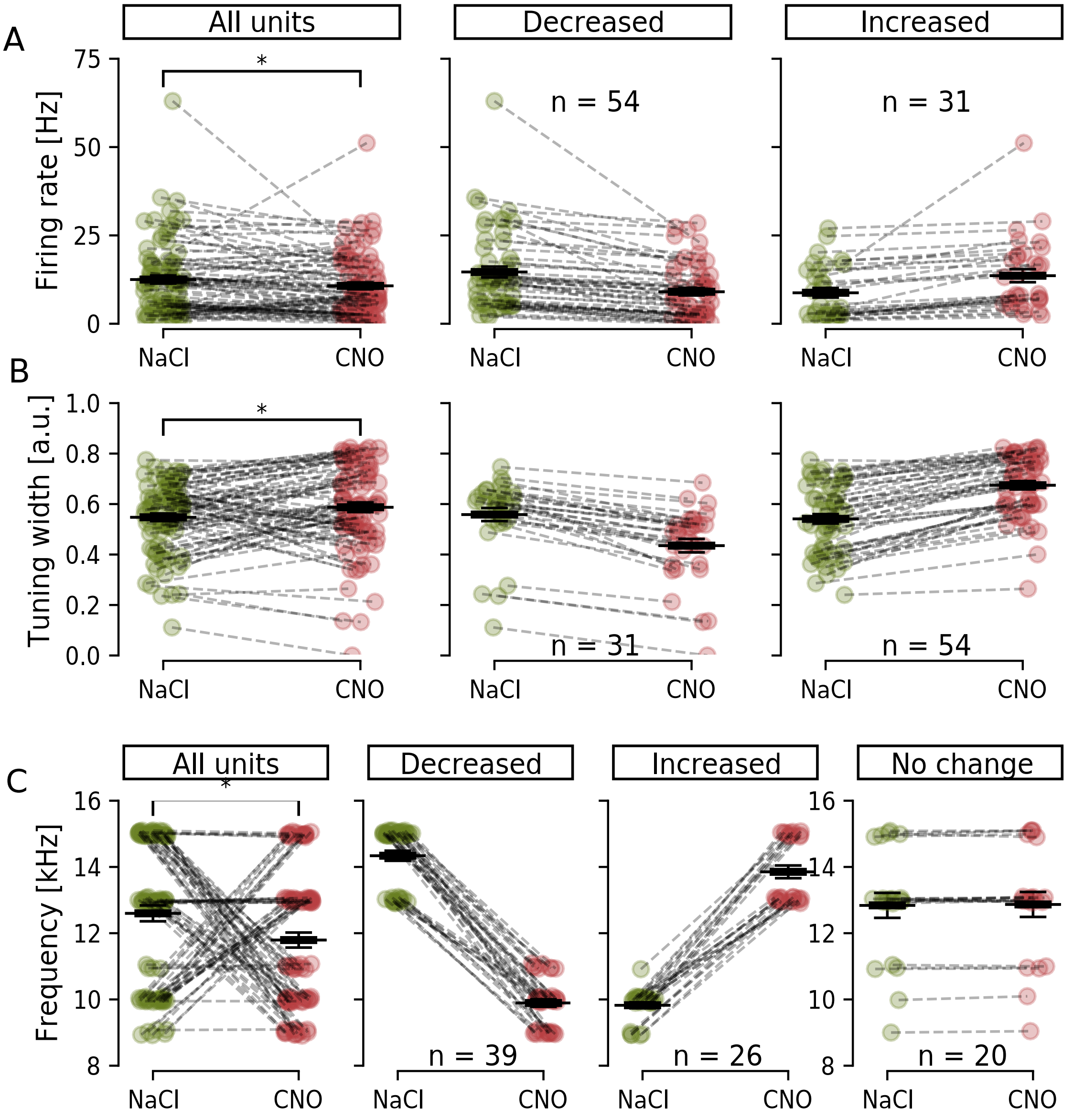


**Supplementary figure S2. Bimodal unit responses seen upon CNO administration in hM4Di+ animals that were treated with CNO also during noise exposure**. A) Left; Firing rate in response to 80dBSPL at best frequency for all units (n = 85) from hM4Di+ mice in response to NaCl or CNO. Middle; Only units decreasing (n = 54) firing rate upon CNO administration. Right; Units increasing (n = 31) firing rate after CNO administration. B) Same as 'A' but for Tuning width, with units decreasing (n = 31) and increasing (n = 54) tuning with after CNO administration. C) Same as 'A' for representation of Best frequency in kHz, with units decreasing (n = 39), increasing (n = 26) or maintaining (n = 20) Best frequency response upon CNO administration. Note that units responding to sound do not need to be CaMKIIα+, the unit altered firing properties are in response to sound when CNO is decreasing activity of CaMKIIα+ units of the DCN circuit. *: $p < 0.05$
